# Pharmacokinetics, Tissue Distribution, and Formulation Study of a Small-molecule Inhibitor of MKLP2, LG157

**DOI:** 10.1101/2023.12.25.573327

**Authors:** Namrta Choudhry, Jinhua Li, Ting Zhang, Jiadai Chenyu, Xiaohu Zhou, Qiong Shi, Gang Lv, Dun Yang

**Affiliations:** Chengdu Anticancer Bioscience, Chengdu 610000, China; J. Michael Bishop Institute of Cancer Research, Chengdu 610000, China

**Author notes:** Correspondence authors, (Namrta Choudhry) (Jinhua Li). N.C. and J.L. contributed equally to this paper. Lead contact (Namrta Choudhry).

**Keywords:** **LG157**, MKLP2, pharmacokinetics, tissue distribution, preclinical formulation, SEDDS

## Abstract

**LG157** is a recently identified small-molecule inhibitor of mitotic kinesin-like protein 2 (MKLP2), an overlooked oncology target. This study aims to explore the drug developability of **LG157**, by assessing its druglike properties, determining plasma drug exposure in various oral formulations, and exploring the self-emulsifying drug delivery system (SEDDS). Solubility of **LG157** ranges from 175 to 228 μM across pH 1.0 to 13.0, with a Log*D* of 2.41 at pH 7.4. It showed a high protein binding rate of 92.58% in mouse plasma and 90.30% in human serum. The bioavailability radar plot aligns with experimental data (69-85%), indicating good bioavailability. In line with the computation prediction, preclinical formulation studies in mice reveal that all five formulations tested offer decent plasma **LG157** exposure, with the highest level of **LG157** exposure in the PEG300-based formulation. Subsequent tissue distribution studies in rats indicated that the compound is widely distributed with the highest concentration of **LG157** in the liver and the lowest level in the brain. The optimal SEDDS formulation, SEDDS-F14, consists of 65% Oleic acid, 26.25% Tween 20, and 8.75% PEG400 as oil, surfactant, and co-surfactant, respectively. SEDDS optimization, based on the central composite design, has achieved the maximum loading of 188.7 mg/mL for **LG157**. These findings support the developability of **LG157** and encourage continued exploration and refinement of formulations for improved therapeutic efficacy.

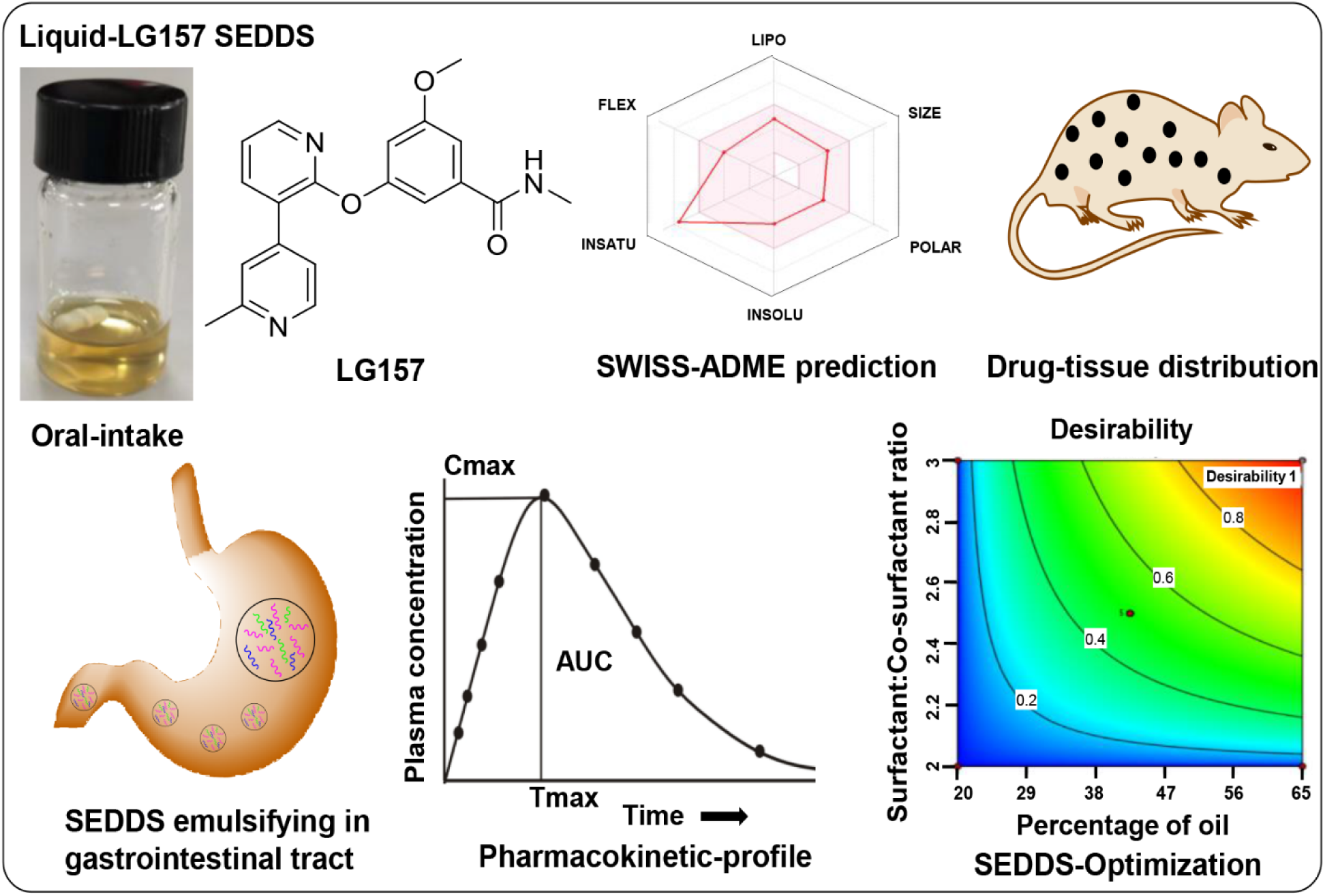

## INTRODUCTION

Cancer remains a formidable global health challenge, with various malignancies contributing significantly to worldwide morbidity and mortality.^1^ Central to the complexity of cancer is the MYC oncoprotein, a major oncogenic driver overexpressed in up to 70% of human cancers.^2^ Overexpression of MYC is linked to poorly differentiated, aggressively progressing tumors. Despite considerable research into MYC biology, this oncoprotein is still largely “undruggable,” with no targeted therapies effectively countering it.^3^ The challenges in directly targeting MYC are compounded by its critical roles in diverse physiological functions, raising concerns about potential severe adverse effects if MYC is disabled in normal tissues.

In the landscape of cancer genome-informed treatment, synthetic lethal strategies have emerged as a promising alternative to oncoprotein-targeted cancer therapies.^4^ **LG157** is a potent, orally available small molecule that targets the mitotic kinesin-like protein 2 (MKLP2)^5^, also termed kinesin family member 20A (KIF20A).^6^ **LG157** is an amide group derivative named 14a (2-methyl pyridine) which has been reported for delocalizing Aurora kinase B, in our recently published study.^7^ Disablement of this underutilized target by **LG157** has been shown to elicit a synthetic lethal interaction with MYC. It selectively eliminates MYC-overexpressing cells while sparing normal dividing cells.^7^ The therapeutic potential of **LG157** is validated through growth suppression in nearly 100 human cancer cell lines *in vitro* and notable tumor growth inhibition in mice.

The promising attributes of **LG157** necessitate detailed investigations into its solubility, stability, physicochemical properties, pharmacokinetics (PK), formulation, and, therefore, guiding further lead optimization or enabling a smooth future transition to clinical trials. This information is crucial for determining the compound’s safety, efficacy, developability, and dosage guidelines. While various conventional oral drug delivery methods, such as salt formation and nanocrystal technology, have been successful, the last few years have seen the emergence of self-emulsifying drug delivery systems (SEDDS) as an advanced method offering unique advantages.^8^

SEDDS, a sophisticated drug delivery approach for both hydrophobic and hydrophilic drugs, has the potential to address the poor and inconsistent bioavailability issues inherent in some traditional formulations.^9^ SEDDS is the composition of an isotropic mixture of oil, surfactant, and co-solvent. Upon contact with body fluids, it spontaneously forms nanodroplets that rapidly spread and enhance self-emulsification in the gastrointestinal tract (GIT).^10^ This drug delivery system could enhance the solubility of the drug in the GIT and circumvent hepatic first-pass metabolism.^11^

SEDDS fundamentally differs from conventional oral drug delivery systems by utilizing enzymatic digestion to release drugs in the GI tract.^12^ This unique mechanism of SEDDS like other lipid-based delivery methods offers several advantages: protection before absorption, inhibition of P-glycoprotein-mediated (PGP) drug efflux, increased permeability and lymphatic transport, controlled plasma drug concentration profiles, improved drug loading efficiency, and enhanced bioavailability.^8^

This study aims to investigate **LG157** regarding physiochemical properties, druglike properties, PK properties *in vivo*, and tissue drug distribution to assess its development potential as an oral drug. **LG157** is classified as a Class I drug in the Biopharmaceutics Classification System (BSC), indicating ease of formulation with conventional methods. Strategically, we also explored SEDDS for **LG157** in this study because this formulation represents an opportunity to transcend the limitations of standard methods, potentially enhancing **LG157**’s bioavailability and therapeutic efficacy. Demonstration of the feasibility of formulating **LG157**-SEDDS opens an avenue for future exploration in this direction.

## EXPERIMENTAL SECTION

### Solubility and Stability Assays

#### *In vitro* Stability in Plasma and Serum

Human serum AB (Cat. #100512) was sourced from Geminibio in California, USA, and mouse plasma (Cat. #SND-X0102) was obtained from Shinuoda in Chuzhou, China. The *in vitro* stability of **LG157** was examined at concentrations of 1 µM and 10 µM. To prepare the samples, blank mice and human serum were spiked with the compound, diluted from a stock solution with acetonitrile, in a 1.5 mL Eppendorf tube (ensuring that the organic solvent content was less than 2.5%). These samples were then incubated for 2 h at 37.5 °C in a water bath. Subsequently, 40 μL of the incubated samples were extracted with 460 μL of acetonitrile by vortexing at 3000 rpm for 2 minutes and then centrifuged at 14,000 rpm for 10 minutes. Finally, 150 μL of the supernatant from each Eppendorf tube was transferred into the liner pipeline of 1.5 mL vials for analysis by Liquid Chromatography-Mass Spectrometry (LC-MS).

#### pH Stability and Solubility

An aliquot of 10 μL of **LG157** was diluted with 990 μL of pH buffer (pH: 1.0 (HCl); 2.2 (HCl); 4.6 (0.01 M citrate buffer); 6.0 (0.01 M PB); 6.8 (0.01 M PB); 7.4 (0.01 m PBS); 8.0 (0.01 M Tris-HCl); 9.0 (0.01 M Tris-HCl); 13.0 (0.01 M NaOH)) in 1.5 mL Eppendorf tubes, resulting in samples with concentrations of 200, 100, and 50 μM. At each specified time, the aliquots of 100 μL were removed from the tubes and then mixed with 100 μL acetonitrile to quantify **LG157** and to detect any degradation products using LC-DAD-MS.

#### Humidity, Strong Light, and Thermal Stability

Stability assays of **LG157** are estimated by drug stability test chamber and drug photostability chamber that are discussed in the International Conference on Harmonization (ICH) guidance.^13^ **LG157** was subjected to LC-MS for purity and stability after chamber placement duration.

#### Octanol-PBS Partition Coefficient (Log*D*)

The octanol phase was thoroughly mixed at a 1:1 volume ratio (Vol/Vol: 50/50) with an aqueous solution (either HCl, pH 2.2, or PBS, pH 7.4 buffer), and the mixture was allowed to saturate for 24 h. Subsequently, 2 μL of the **LG157** stock solution (20 mM, dissolved in DMSO) was added to 0.8 mL of this saturated mixture in a 1.5 mL Eppendorf tube. The tube was then shaken at 350 rpm for 24 h at room temperature. Following this, the aqueous and organic phases were permitted to separate over one hour. Both phases were subsequently quantified for **LG157** using LC-MS.

The logarithm of the distribution coefficient (Log*D*) was calculated using the following equation:

Log*D* octanol/water= Log (Concentration in octanol/Concentration in water).

#### Plasma and Serum Protein Binding

Equilibrium dialysis is considered a reference method to evaluate plasma protein binding with modification.^14^ The samples (10 μM) were prepared by spiking blank mouse and human serum with **LG157**, diluted from a stock solution with acetonitrile (ensuring that the organic solvent content was less than 2.5%). The samples were then incubated in a water bath for 2 h at 37.5 °C to allow for equilibrium of binding interaction. Following this, 200 μL of the protein solution was dialyzed for 18 h at 37°C against 40 mL of dialysis buffer (0.01M PBS, pH 7.4) using snakeskin dialysis tubing (Thermo Scientific, 10 K MWCO). Finally, the dialysis buffer was collected and analyzed to quantify the **LG157** by LC-MS.

The plasma/serum protein binding rate was calculated using the following equation:

Binding rate (%) = (Conc. plasma chamber – Conc. buffer chamber)/Conc. plasma chamber × 100%

#### LC-MS (or LC-DAD-MS) Conditions

The LC-MS analysis was performed on an LC-MS system (Agilent InfinityLab LC/MSD iQ) with a Poroshell 120 SB-C18 column, 2.1 × 50 mm, 2.7 μm (Agilent, California, USA). Flow rate, 0.5 mL/min; column temperature, 25°C. The mobile phase comprised pure water supplemented with 5 mM NH_4_HCO_3_ (phase A) and HPLC-grade methanol (phase B). The gradient system used: 0 - 3.5 min, 20 → 95% B; 3.51 – 5.0 min, 95% B; 5.01 – 6.5 min, 20% B at the flow rate of 0.5 mL/min. The injection volume: 2 µL; DAD detection was set at a wavelength of 254 nm. MS conditions: ion source, ESI; scan polarity, positive ion; drying gas, nitrogen gas at the flow rate of 13 L / min; temperature, 325 ◦C; ion spray voltage, + 3500 V; collision energy, 100 V; spraying gas (N2) pressure: 55 psi; monitor mode, selected ion monitoring (SIM). DAD 254 nm or pseudomolecular ion ([M+H] +), m/z 350.2 for **LG157** were detected for quantitative analysis.

#### Computational analysis of Lipinski parameters and drug-likeness properties

Pharmacokinetics and drug-likeness prediction of **LG157** were performed by the online tool Swiss ADME predictor software at the Swiss Institute of Bioinformatics.^15^ Various Lipinski parameters included (MW, molecular weight; HBA, Hbond acceptor; HBD, H-bond donor; TPSA, topological polar surface area; cLogP, *n*-octanol/water distribution coefficient), and the number of rotatable bonds (nrotb) were calculated using Swiss-ADME.

### Preclinical Formulation Preparation and PK studies

#### 50% PEG 300 Formulation

Prepare a 10% Tween 80 solution by mixing 10 mL of Tween 80 with 90 mL of ddH_2_O, then vortex and place in a water bath ultrasonic for 20 minutes to achieve a uniform solution. For the PEG 300 mixture, combine 50 mL of PEG 300 with 28 mL of ddH_2_O and swirl on a vortex mixer, followed by adding 10 mL of the 10% Tween 80 solution, and vortex again. The resulting pH will be approximately 5.6, which should be reduced to 2.2 using methane-sulphonic acid. Adjust the volume to 90 mL with ddH_2_O to finalize the PEG 300 mixture. For the **LG157** solution, dissolve the solid powder into DMSO by vortexing, followed by 10 minutes of sonication in a water bath ultrasonic (Bransonic 1510). Then, dilute the **LG157** solution at a ratio of 1:9 (v/v) with the PEG 300 mixture, blending by vortexing and sonicating for an additional 10 minutes in the water bath ultrasonic to yield a completely dissolved clear solution.

#### 100% PEG400 Formulation

Powder of **LG157** was completely dissolved in 100% PEG 400 by vortexing and sonicating for 10-30 minutes in a water bath ultrasonic.

#### 10% DMA (pH 2.2) Formulation

Add 15 mL of EtOH to a beaker containing 10 mL of N, N-Dimethylacetamide (DMA), and stir until well blended. Next, add 30 mL of 1,2-Propanediol (PG], stirring until thoroughly mixed, followed by the addition of 40 mL of ddH_2_O. Adjust the pH to 2.2 using diluted HCl, and then add ddH_2_O to make up the total volume of the mixture to 100 mL. To prepare the **LG157** solution, dissolve the powder of **LG157** in the 10% DMA solution by vortexing and sonicating for 10-30 minutes in a Bransonic 1510 water bath ultrasonic.

#### 0.5% HPMC Formulation

Begin by adding 10 mL of Tween 80 to 90 mL of ddH_2_O, then vortex the mixture and place it in a Bransonic 1510 water bath ultrasonic for 20 minutes to create a uniform 10% Tween 80 solution. Next, transfer 80 mL of ddH_2_O into a 200 mL beaker equipped with a magnetic stirrer. Slowly add either 0.5 g of Hydroxy Propyl Methyl Cellulose (HPMC) powder to the beaker, stirring continuously until the powder is completely dissolved. Add 10 mL of the 10% Tween 80 solution to the beaker, and after adjusting the pH to the range of 1-2.2 using concentrated HCl, make up the volume to 100 mL. Finally, add **LG157** powder to the HPMC solution and homogenize for 30 minutes using an ultrasonic cell disruptor (Ning Bo Xin Zhi, JY92-IIN), to produce a milky white uniform suspension.

#### 100% oleic acid Formulation

The powder of **LG157** was completely dissolved in 100% oleic acid by vortexing and sonicating for 10-30 minutes in a Bransonic 1510 water bath ultrasonic.

All the following **LG157** formulations should be stored at 4°C and stable for at least one month.

#### Ethics statement

The study was conducted according to the animal study protocols with a reference number (BUA#: BU000021-010) and was approved by the institutional animal care and use committee (IACUC guideline 20.20) of the Institutional Biosafety Committee (IBC) (or ethical committee) at the J. Michael Bishop Institute of Cancer Research (MBICR). Directions for CO_2_ euthanasia of rodents (mice) were done under (IACUC guideline 20.20). This study was carried out in compliance with the ARRIVE guidelines.^16^

#### Rapid PK Assay

For PK and tissue distribution analysis, unless otherwise described, the animals were starved for 16 h before drug administration, with food being allowed 2 h post-treatment. Free water intake was permitted throughout the study. The samples from blood and tissues were quantified for **LG157** using LC-MS/MS.

#### PK of LG157 in Mice

For the PK analysis of **LG157** in various formulations, two different groups of mice were used. For the 50% PEG 300, HPMC, and DMA formulations, two 11-week-old female FVB/N mice, weighing between 22-24 g each, were used. For the 100% PEG 400 and oleic acid formulations, three 21-week-old female BAL B/c mice, weighing between 18-23 g each, were employed. Each mouse received an oral dose of 50 mg/kg of **LG157**. Blood samples (100 μL) were collected into heparinized tubes at time intervals of 0, 0.5, 1, 3, or 6 h post-drug administration.

#### Drug Tissue Distribution in Rats

Nine 9-week-old female Wistar rats, weighing between 200-240 g each, were orally administered 50 mg/kg of **LG157**, dissolved in 50% PEG 300 at pH 2.2. At intervals of 1, 4, and 8 h post-administration, orbital blood of the rat was firstly collected, and then 1x PBS was used for systemic perfusion to expel blood from the tissues, finally various organs including heart, lung, liver, pancreas, spleen, ovary, kidney, leg muscle, lymph nodes, adipose tissue, and brain were collected. Tissues and organs were carefully washed with 1 x PBS to remove residual blood on the surface, and approximately 200 mg of tissues were combined with 800 μL of grinding fluid (methanol: PBS = 1:1), homogenized, and centrifuged at 12000 rpm for 5 minutes. The supernatant was utilized for the quantification of **LG157**. The plasma **LG157** concentration was also analyzed using LC-MS/MS.

### Determination of the Drug Concentration in Plasma and Tissues

#### LC-MS/MS Conditions

The drug concentration in plasma and tumor tissues was determined using a Thermo Scientific Vanquish UHPLC system coupled with a TSQ Quantis™ Triple Quadrupole Mass Spectrometer, equipped with a heated-electrospray ionization (H-ESI) source (Thermo Scientific, Santa Clara, California, U.S.A.). System control and data processing were managed through Chromeleon Console Chromatography Data System (CDS) software (Version 7.2.10 ES, Thermo Fisher, San Jose, USA). The chromatographic conditions included an ACQUITY UPLC BEH C18 column (2.1 mm × 50 mm, 1.7 μm particle size) (Waters, MO, USA) at 25 °C, and the mobile phase consisted of A, 5 mM NH4HCO3 in water, and B, acetonitrile. Gradient elution was performed as follows: 0 - 0.5 min, 10% B; 0.5 - 3 min, 10 → 95% B; 3 - 4 min, 95% B; 4.01 - 5.5 min, 10% B; at a flow rate of 0.3 mL/min. Samples were kept at 6 °C in the automatic sampler, and the injection volume was set at 5 μL. Mass spectrometric analysis was acquired in the positive ion mode with nitrogen as sheath and auxiliary gas and argon as the collision gas, with the following main parameters: spray voltage, 3500 V; sheath gas pressure, 35 Arbitrary units (Arb); auxiliary gas pressure, 10 Arb; collision gas pressure, 1.5 mTorr; ion transfer tube temperature, 320 °C; vaporizer temperature, 350 °C. Quantification was performed in the SRM mode. The RF LENS and source CID for **LG157** were set as 131.2 V and 18.4, respectively. The SRM transitions were m/z 350.20 > 319.0 (quantifier) (collision energy 20 V) and m/z 350.20 > 276.0 (qualifier) (collision energy 34 V).

#### Sample Preparation and Processing

**LG157** in the tissue fluid and plasma samples was analyzed quantitatively using LC-MS/MS. Calibration standards, ranging from 0.025 to 12.5 μM, and quality control (QC) standards at three levels (0.075, 4, 9 μM), were prepared by spiking blank plasma and tissue fluid with the test compound. A 10 μL aliquot of either plasma or tissue fluid sample was mixed with 10 μL of the internal standard work solution in a 0.5 mL Eppendorf tube, followed by extraction with 200 μL of acetonitrile. The mixture was vortexed for 5 minutes and centrifuged for 15 minutes at 14000 rpm at room temperature. The resulting supernatant was then analyzed using the aforementioned LC-MS/MS system.

### LG157-SEDDS Design and Characterization

#### Excipients Source

Various oils, surfactants, co-surfactants, and polymers were employed to assess the solubility of **LG157**. The oils used in the study included Oleic Acid (Cat. #BD00781195 Bidepharm), Castor Oil (Cat. #C110663 Aladdin) CCT, Caprylic Capric Triglyceride (CCTG) (Cat. #H860153 Macklin), Isopropyl Palmitate (IPM), Olive Oil (Cat. #O108686 Aladdin), and Soybean Oil (Cat. #S110245 Aladdin). The surfactants comprised Labrasol (Cat. #190092 Gattefosse), Polysorbate-80 (Tween-20) (Cat. #T104863 Aladdin), Polysorbate-80 (Tween-80) (Cat. #T104866 Aladdin), Ethoxylated hydrogenated castor oil (Cremophor RH40) (Cat. #E875014 Macklin), and Castor oil ethoxylated (Cremophor EL) (Cat. # 70726A Adamas). Co-surfactants in the analysis featured Macrogol 400 (PEG-400) (Cat. #BD00777172 Bidepharm), Macrogol 200 (PEG-300) (Cat. #p103728 Aladdin), Macrogol 200 (PEG-200) (Cat. #20805 MELOPEG), Transcutol HP (Cat. #189790 Gattefosse), Propylene Glycol Monocaprylate NF (Capryol-90) (Cat. #191178 Gattefosse), Span-20 (Cat. #S102859 Aladdin), Span-80 (Cat. #S110840 Aladdin), Lauroglycol-90 (Cat. #D54137 Gattefosse), and Polypropylene Glycol (PPG). Polymers utilized consisted of 10% Polyvinypyrrolidone K30 (PVP-K30) (Cat. #BD148615 Bidepharm), 10% Poloxamer 407(Cat. #R051361 Bidepharm), 10% D-Sorbitol (Cat. #D856745 Macklin), 10% Poloxamer 188, 1% Hydroxypropylmethylcellulose (HPMC) (Cat. #H108820 Aladdin), and 5% Polyvinyl alcohol (PVA-124) (Cat. #P909857 Macklin). All chemicals used here were of analytical reagent grade and were sourced locally.

#### Solubility of LG157 in Various Excipients

A standardized 150 mg amount of **LG157** was mixed with 1g of an excipient in a sealed vial and stirred at 500 RPM at 37°C for 72 h. If the solution became clear, indicating complete dissolution, an additional 50 mg of **LG157** was added and incubated. The solubility of **LG157** in each excipient reached saturation when the mixture appeared slightly turbid. Samples were then centrifuged at 14,000 rpm for 20 minutes, and around 20-25 mg of the supernatant was weighed, collected, diluted with DMSO, and analyzed using LC-DAD-MS (the conditions are the same as the above LC-DAD-MS method). A linear calibration curve for **LG157**, with concentrations ranging from 0.015625 to 1mg/ml and correlation coefficients above 0.999, was established.

#### Construction of a Pseudo-ternary Phase Diagram

Oleic acid, Tween 80, and PEG400 were employed as the oil phase, surfactant, and co-surfactant, respectively, for constructing a pseudo-ternary phase diagram using the water titration method described previously.^17^ To identify the optimal ratios of these three components in microemulsions, S_mix_ (surfactant: co-surfactant mixture) compositions at a 3:1 w/w ratio were used. Predefined w/w ratios of oil to Smix (7:3, 6:4, 5:5, 4:6, 3:7, 2:8, and 1:9) were mixed and 10 µl of the mixture was titrated into 100 mL of water at 37°C with magnetic stirring at 10 rpm. The emulsification tendency was assessed visually for turbidity and viscosity.

When oil droplets readily coalesced into a fine, milky emulsion and spread readily in water, the emulsion was considered ‘good’. When there was poor or no emulsion formation with immediate coalescence of oil droplets, especially after stirring was stopped, it was considered ‘bad’. The corresponding phase diagrams delineated the boundaries between the surfactant, co-surfactant, and oil phases, using an emulsion-dotted area serving as the index for phase suitability and component proportion. The optimal ratio of oil to Smix (6.5:3.5 w/w) was identified and fixed by comparing the areas of the emulsion regions on the phase diagrams, using Origin Pro 8.0 software for visualization. The optimal ratio (6.5:3.5 w/w) for **LG157** SEDDS, categorizing it as class II SEDDS. This class is characterized by high oil content and water-insoluble surfactants, based on the lipid formulation classification system by Pouton^18^.

#### Optimization of the LG157-SEDDS Formulation

The formulated SEDDS for **LG157** incorporated optimal excipients identified through solubility studies and pseudo-ternary phase diagrams, with a preferred oil-to-surfactant/co-surfactant ratio of 6.5:3.5. The solubilization potential and extensive emulsification region in the phase diagram are the major factors that guide the selection of the relative amount of SEDDS components. The optimization of the formulation composition is a complex process that requires extensive experiments. To further refine this formulation, Central Composite Design-Response Surface Methodology (CCD-RSM) was employed^19^, using drug loading as the response variable. CCD-RSM enabled the exploration of multiple variables across various levels with minimal experimental runs.^20^

#### Design of Experiments (DoE)

A two-factor, two-level face-centered CCD-RSM was employed for the formulation of SEDDS containing **LG157**. This approach aimed to assess the influence of individual excipient concentrations and their interactions on **LG157** loading within the SEDDS. This study utilized two factors, Factor A (X1) which represented the oil concentration, and Factor B (X2) which represented the ratio of surfactant to co-surfactant (S_mix_) to investigate their influence on drug loading capacity (Y1, mg/mL). The effects of these formulation components (Factors A and B) on the variable (Y1: **LG157** loading) were assessed using the Design of Expert (DOE) software. Drug loading capacity is an important parameter to be considered while selecting the oil, surfactant, and cosurfactant. A total of 13 experimental designs were generated by the software and performed in a randomized sequence to enhance predictive accuracy. To increase the predictability, experiments were performed in random order. The collected data were analyzed using CCD-RSM within the same software to evaluate the optimized value for **LG157** loading in the SEDDS.^20^ The CCD-RSM was designed using Design of Expert (DOE) 8.0.4.1 software (MN, USA) and executed randomly.

#### Characterization and Evaluation of LG157-SEDDS

The **LG157**-SEDDS formulations were subject to a visual assessment for clarity, homogeneity, and color. Screw-cap vials containing the formulations were inspected under light for clarity, while color was assessed via normal visual examination. The presence or absence of precipitation was observed to gauge formulation homogeneity. These formulations were subsequently stored at room temperature for 15 days for further visual assessments.

**a. Self-Emulsification Time.** All the formulations were evaluated for emulsification time as reported.^21^ One milliliter of optimized formulation was mixed with 200 ml of distilled water under agitation at 100 rpm with a magnetic stirrer. The time required for emulsification was recorded.
**b. Precipitation Assessment.** Precipitation assessment was done by visual inspection of the resultant emulsion after storage for 24 h at room temperature. The formulations were classified as clear (transparent or transparent with a bluish tinge) non-clear (turbid), stable (no precipitation at the end of 24 h), or unstable (showing precipitation within 24 h).^22^ All studies were performed in triplicates.
**c. Transmittance Percentage**. The percent transmittance of diluted samples was determined using a UV spectrophotometer at 620 nm against a reference blank solution.^23^

## RESULTS

### Solubility, Stability, Log*D*, and Plasma Protein Binding

**LG157** exhibited high aqueous solubility ranging from 175 to 228 μM from pH1.0 to pH13.0 (**Figure 1A**). It demonstrated exceptional pH stability under the acidic condition of the stomach (pH 2.2), as well as under the physiological condition (pH 7.4) for up to 72 h (**Table 1**). This stability suggests that the activity of **LG157** would remain unaffected even with varying pH levels in the human body.

**Figure 1.**
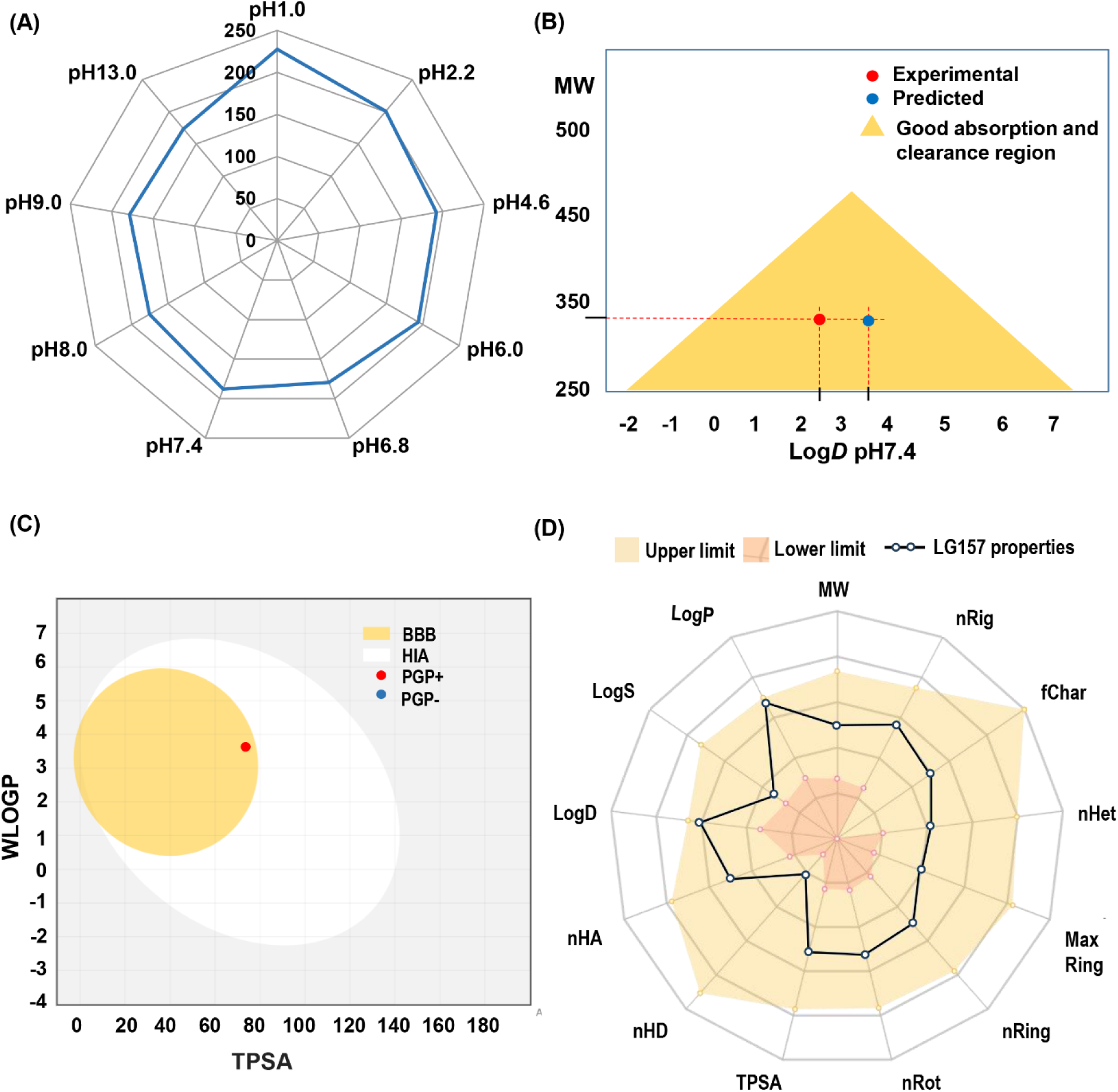
Solubility and physicochemical property analysis of **LG157**. (A) Solubility of LG157 at pH 1.0 to 13.0. (B) Golden triangle plot with predicted and experimentally determined Log*D* at pH 7.4 to estimate *in-vitro* permeability and clearance trends. Y axis, molecular weight (MW). X axis, distribution coefficients (log*D*) of LG157.

**Table 1.**
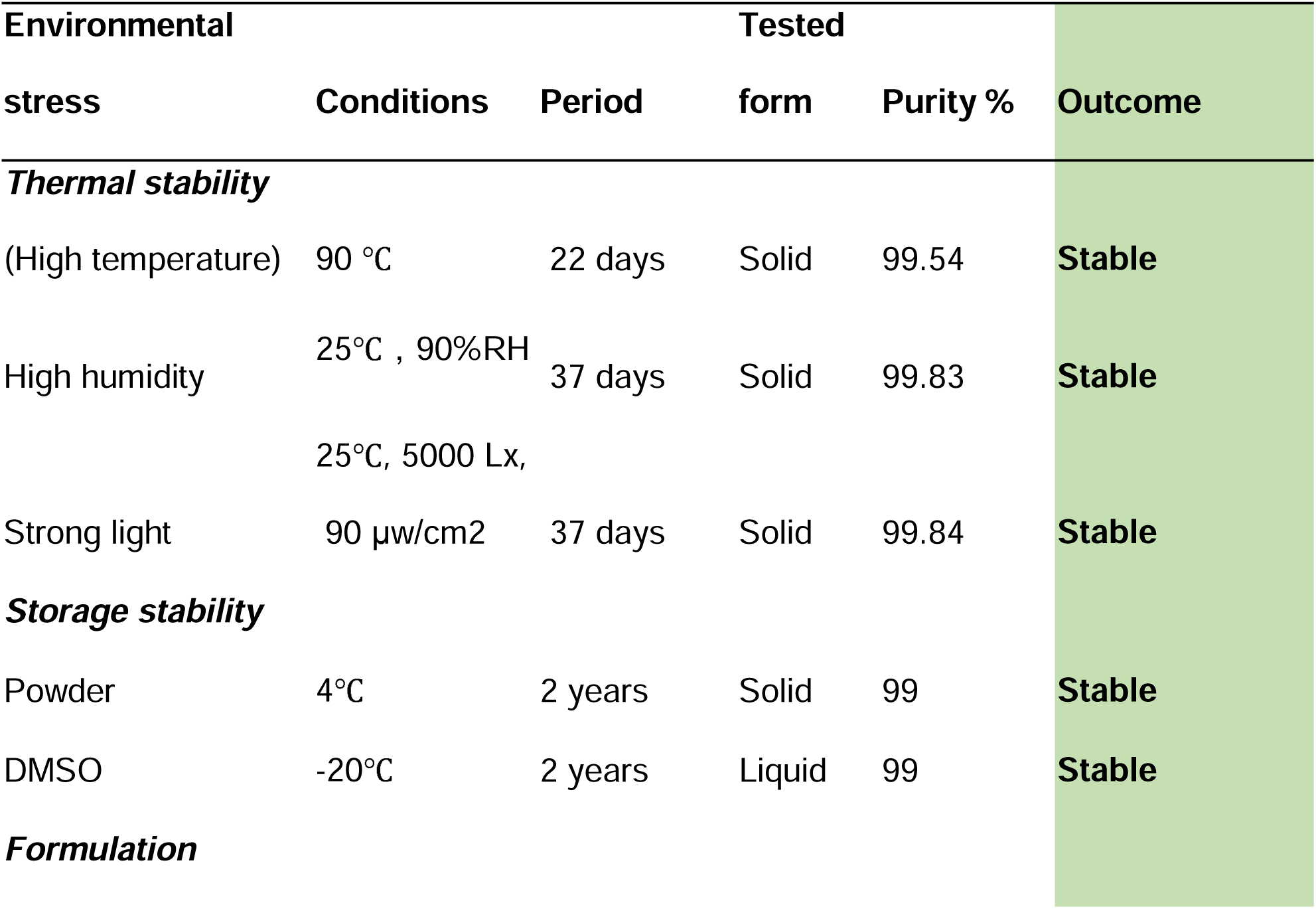
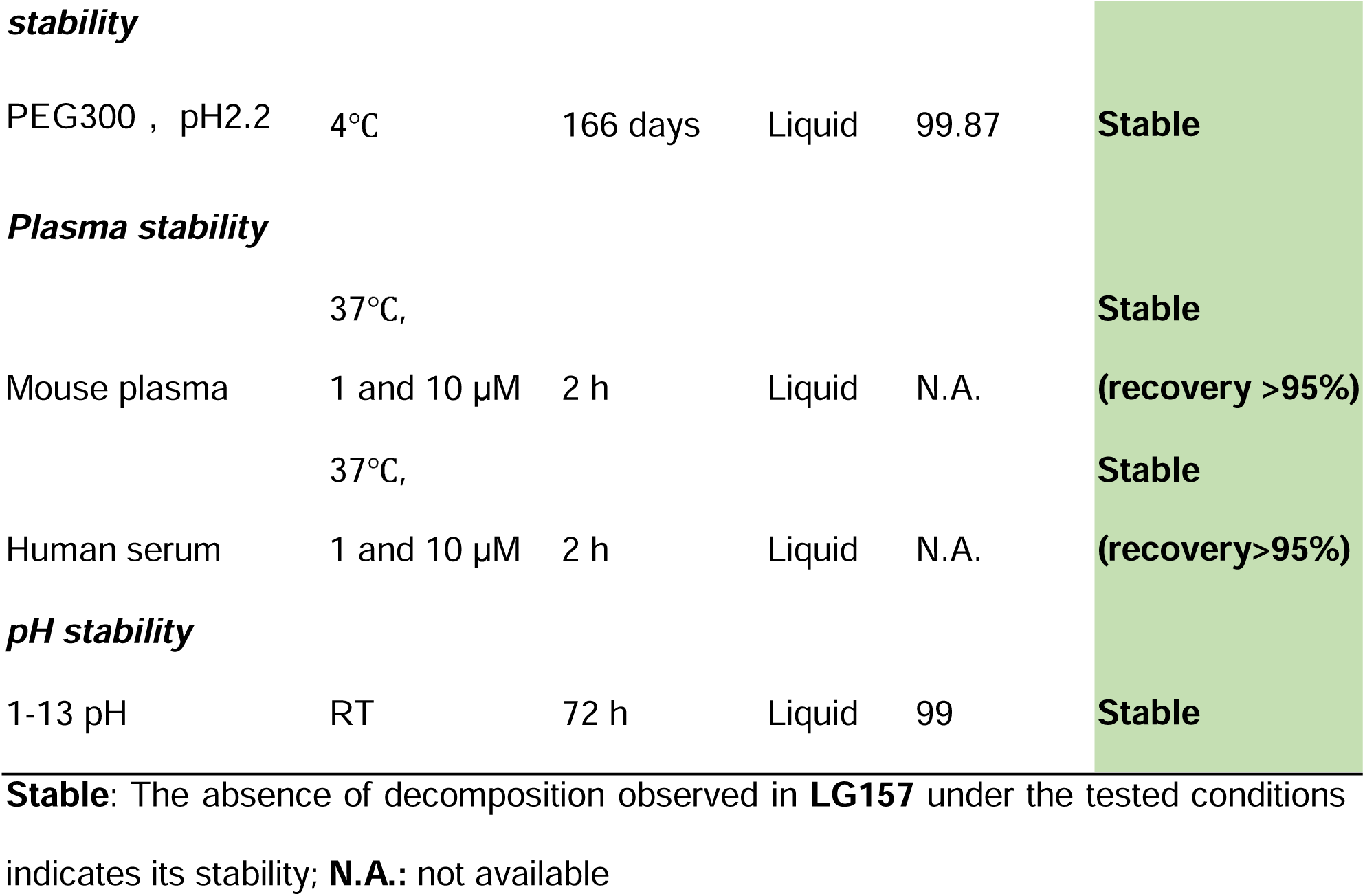
Stability of LG157 Under Different Environmental and Stress Conditions.

| Environmental |  |  | Tested |  | Outcome |
| --- | --- | --- | --- | --- | --- |
| stress | Conditions | Period | form | Purity % |  |
| ***Thermal stability*** |  |  |  |  |  |
| (High temperature) | 90 °C | 22 days | Solid | 99.54 | **Stable** |
| High humidity | 25°C , 90%RH | 37 days | Solid | 99.83 | **Stable** |
|  | 25°C, 5000 Lx, |  |  |  |  |
| Strong light | 90 μw/cm2 | 37 days | Solid | 99.84 | **Stable** |
| ***Storage stability*** |  |  |  |  |  |
| Powder | 4°C | 2 years | Solid | 99 | **Stable** |
| DMSO | -20°C | 2 years | Liquid | 99 | **Stable** |
| ***Formulation*** |  |  |  |  |  |
| **stability** |  |  |  |  |  |
| PEG300 , pH2.2 | 4°C | 166 days | Liquid | 99.87 | **Stable** |
| **Plasma stability** |  |  |  |  |  |
|  | 37°C, |  |  |  | **Stable** |
| Mouse plasma | 1 and 10 µM | 2 h | Liquid | N.A. | **(recovery >95%)** |
|  | 37°C, |  |  |  | **Stable** |
| Human serum | 1 and 10 µM | 2 h | Liquid | N.A. | **(recovery>95%)** |
| **pH stability** |  |  |  |  |  |
| 1-13 pH | RT | 72 h | Liquid | 99 | **Stable** |

**Stable:** The absence of decomposition observed in **LG157** under the tested conditions indicates its stability; **N.A.:** not available

Stability assessments of **LG157** under stress conditions, including high temperature, high humidity, and strong light, for 22-37 days revealed an impressive stability profile (**Table 1**), indicating its storage suitability under potentially challenging conditions. **LG157** displayed a moderate degree of lipophilicity as its Log*D* at pH 7.4 was experimentally determined as 2.41, which enables good cellular permeability. The plasma protein binding fractions of **LG157** were 92.58% in mouse plasma and 90.30% in human serum (**Table 2**). In line with the Biopharmaceutics Classification System (BCS),^24^ **LG157** is categorized in the BCS I. Thus, the **LG157** exhibited a superior stability profile, adequate aqueous solubility, and appropriate lipophilicity, making it a promising candidate for oral drug formulation. Next, we performed a physicochemical and pharmacokinetic investigation of this compound.

**Table 2.** Log*D* and Plasma Protein Binding (PPB) of LG157.

| **LogD** |  | **PPB** |  |
| --- | --- | --- | --- |
| pH2.2 | pH7.4 | Mouse plasma<br>(drug conc.: 10 µM) | Human serum<br>(drug conc.: 10 µM) |
| -0.77 | 2.41 | 92.58±0.43% | 90.30±0.87% |

### ADME predictions for LG157

The idea of “drug-likeness” was put forth to offer helpful suggestions in the early stages of oral drug discovery, increasing the likelihood that a chemical entity will complete the later stages of drug development.^25^ It can be defined as the sum of the molecular physicochemical properties that are characteristics of oral drugs. Indeed, drug-likeness is often referred to as PK properties and safety, or compounds with desirable ADMET (Absorption, distribution, metabolism, excretion, and toxicity) properties.^26^ In the drug development process, permeability, gastrointestinal absorption, and blood−brain barrier (BBB) penetration are critical factors to consider.

The Golden Triangle is a visualization tool that helps to identify metabolically stable, permeable compounds.^27^ The location of **LG157** within the golden triangular-shaped area, with an experimental Log*D* value of 2.41 at pH 7.4, indicates good permeability and low clearance (**Figure 1B**). Likewise, the BOILED-egg plot depicts the possibility of absorption and penetration of small molecules in the gastrointestinal tract and brain as a function of WLOGP (lipophilicity) and TPSA (topological polar surface area) parameters.^28^ **LG175** location in the BOILED-egg plot suggests that the compound can cross the BBB to some extent and is actively effluxed by P-glycoprotein (Pgp) (**Figure 1C**).

The Radar chart provided a 360° snapshot view of the profile using up to 15 properties. The objective is to determine if the test properties fall within qualifying ranges for oral drugs. **Figure 1D** illustrates all physicochemical properties are within the qualifying range. Further, Swiss ADME prediction data are summarized in **Table S1.** Furthermore, **LG157** also fulfills the drug-likeness characteristics defined by the major pharmaceutical companies; Lipinski’s (Pfizer), Veber’s (GSK), and Egan’s (Pharmacia) filters.^29^ The synthetic accessibility score is 3.04, in line with the easy synthesis of **LG157**. There is no alert for Pan assay interfering compounds (PAINS) (**Table S2**).

In summary, an in-silico study for predicting the physicochemical and drug-like properties of **LG157** revealed promising results for the potential of **LG157** as an orally available compound. Therefore, we moved ahead to test the plasma **LG157** exposure in pre-clinical formulations.

Experimental and predicted Log*D* values are indicated by the red and blue dots, respectively. (C) Boiled-Egg plot. The Brain or Intestinal estimated permeation is a function of a compound’s partition coefficient (WLOGP) and topological polar surface area (TPSA). The high probability of human gastrointestinal absorption (HIA), is represented by the egg white region, whereas the yellow region (yolk) represents the high probability of the blood-brain barrier (BBB) penetration. In addition, the red color shows that a test molecule is actively effluxed by P-glycoprotein (Pgp), represented as (PGP+), whereas the blue color implies not a substrate for Pgp, indicated as (PGP−). (D) A radar plot of multiple physicochemical properties of **LG157.** Here the combined shaded area between the upper and lower limit represents the property ranges that are occupied by 95% of approved drugs. The black dotted line represents the properties of the **LG157**. MW: Molecular weight; nRig: Number of rigid bonds; fChar: Formal charge; nHet: Number of heteroatoms; Max Ring: Number of atoms in the biggest ring; nRing: Total number of rings; nRot: Number of rotatable bonds; TPSA: Topological polar surface area; nHD: Number of hydrogen donors; nHA: Number of hydrogen acceptors; Log*D*: Log at physiological pH 7.4; LogS: Log of the aqueous solubility; LogP: Log of the octanol/water partition coefficient.

### Rapid PK profiles of LG157

Exploration of plasma drug exposure across various formulations might provide a robust foundation for subsequent PK and toxicity investigations. FDA-approved excipients, such as Hydroxypropyl Methylcellulose (HPMC), Polyethylene Glycol 300 (PEG 300), and PEG 400, N, N-Dimethylacetamide (DMA), and Oleic Acid formulations were tested for their ability to deliver **LG157** because they vary in their chemical nature. HPMC is a cellulose derivative and a hydrophilic polymer that can encapsulate a wide range of compounds.^30^ PEG300 is a synthetic, non-ionic hydrophilic polymer known for its versatility with various drugs.^31^ DMA, a polar aprotic solvent, is widely used as a drug delivery vehicle.^32^ Oleic acid, a monounsaturated fatty acid, is particularly suited for lipophilic drugs.^33^ Despite their chemical differences, they all serve the common purpose of improving the solubility and bioavailability of even poorly water-soluble drugs, an essential aspect in the formulation of many therapeutic agents. Despite that these formulations all have the shared advantage of enhancing the solubility of poorly water-soluble drugs, the necessity exists to test different formulations to identify the best option arising from their distinct chemical natures and varying interactions with an **LG157**. We compared plasma drug exposure of **LG157** in mice using five different formulations: 50% PEG 300 with pH ranging from 1 to 2.2, 100% PEG 400, 10% DMA with a pH of 2.2, 0.5% HPMC at a pH of 2.2, and 100% Oleic acid. Data are summarized in **Figure 2**. Each of these formulations proved successful, producing adequate plasma drug exposure (**Figure 2B**). This was evidenced by a range of dose-normalized area under the curve (AUC) values from 1.01 ± 0.60 to 2.24 ± 1.45 μM.h/(mg/kg) (**Figure 2C)**. AUC is a key metric for assessing plasma drug exposure and a dose-normalized AUC of 0.1 μM.h/(mg/kg) or higher is correlated with an 80% chance that the oral bioavailability of a test compound is more than 30%.^34^

**Figure 2.**
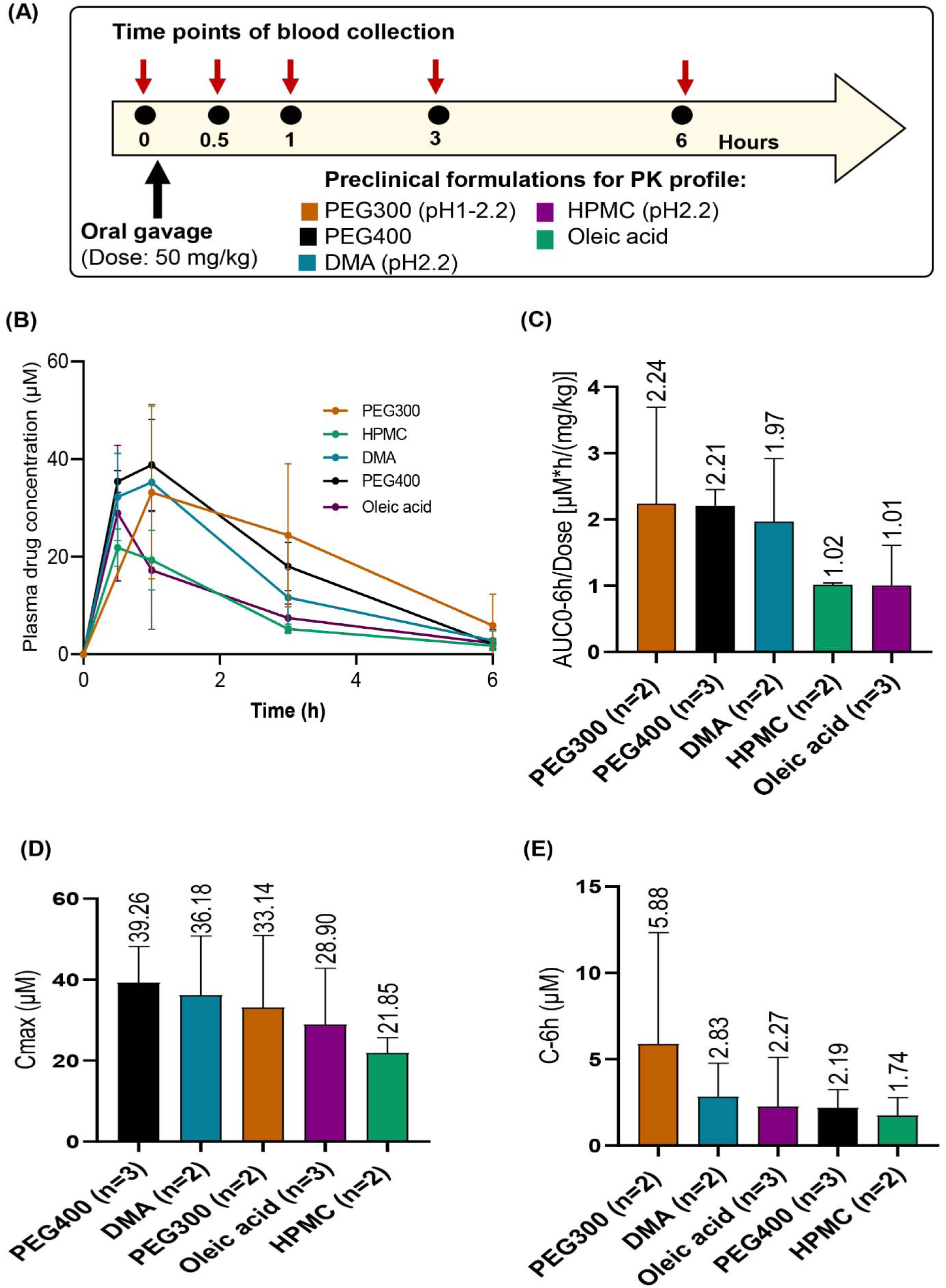
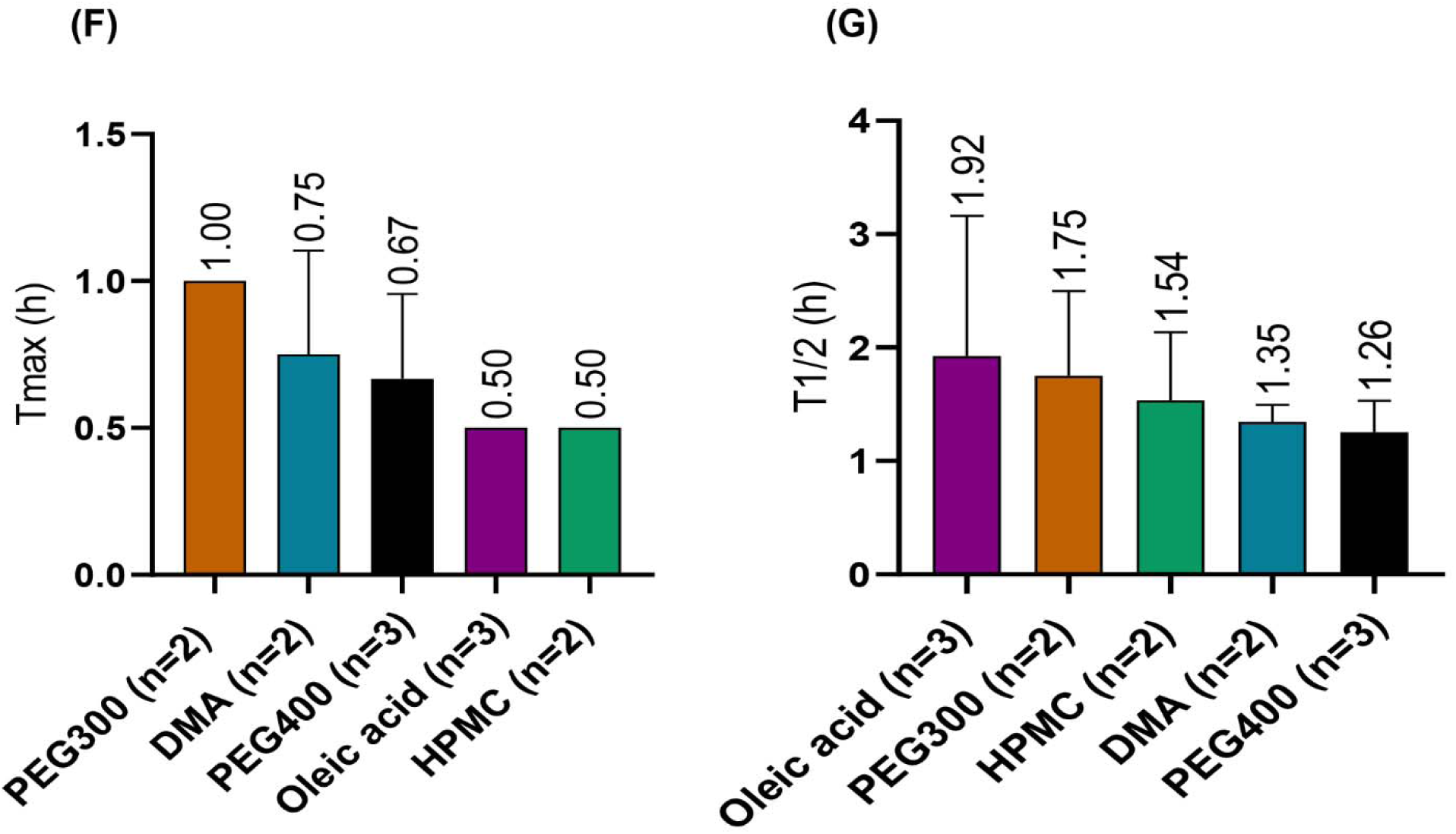
Oral pharmacokinetic parameters of **LG157**. **LG157** was suspended or dissolved into the five types of formulations, including 50% PEG 300, 10% DMA, 0.5% HPMC, 100% Oleic acid, and 100% PEG 400 before being delivered to female FVB/N or BAL B/c) mice. n: the number of mice used in each group. (A) Schematic diagram of timeline for blood collection after a single oral administration of 50 mg/kg of LG157 in the indicated formulation. (B) plasma drug concentration-time curves. (C) The dose-normalized area under the plasma drug concentration-time curve from time zero to 6 h (AUC_0-6h_). (D) The maximum drug concentration in plasma (C_max_). (E) Plasma drug concentration at 6h (C_6h_). (F) Time of peak plasma drug concentration (T_max_). (G) Half-life (T_1/2_). The colors in B-G panels correspond to different formulations described in (A).

Moreover, the peak plasma concentration (C_max_) values ranged from 21.85 ± 3.82 to 39.26 ± 8.91 μM, further highlighting the high plasma drug exposure across different formulations (**Figure 2D**). Based on the mouse plasma protein binding rate of 92.58% for **LG157** in mice (**Table 2**), the plasma peak free drug concentration of 1.63-2.91 μM was inferred and corresponded a 54-97-fold higher than the IC_50_ of 30 nM. This suggests that the drug concentration in the plasma was much higher than the effective concentration required for the target inhibition. Notably, among all the formulations tested, the 50% PEG300 formulation exhibited the highest AUC and drug plasma concentration at six hours (C_6h_), implying that it led to the most sustained plasma drug exposure (**Figure 2E**), In addition, **LG157** in PEG300 formulation exhibited longer T_max_ and a comparable half-life (T_1/2_) relative to **LG157** in Oleic acid formulation (**Figure 2F-G**). As a result, 50% PEG300 was selected as the formulation of choice for further *in vivo* experimentation.

### Oral PK properties of LG157 in rats

PK studies in mice revealed favorable plasma drug concentrations and half-life characteristics for **LG157** (**Figure 2**). We further explored a full PK study of **LG157** in rats by administering the compound via intravenous (IV) injection at 2 mg/kg and orally (PO) at doses of 2, 10, and 50 mg/kg. A dose-proportional increase in both the maximum plasma concentration (C_max_) and the area under the curve (AUC) was observed. Key PK parameters like volume of distribution (V_z_), mean residence time (MRT), and half-life (T_1/2_) also increased in a dose-dependent manner, particularly when escalating from 2 to 10 mg/kg (**Table 3**). The **LG157** showed good bioavailability, ranging 69-85% across the dosages, and displayed no impact on the general well-being of rats up to 50 mg/kg. At the 50 mg/kg oral dose of **LG157**, C_max_ reached 34.66 ± 6.48 μM, AUC_0-∞_ was 114.11 ± 20.94 μM·h, T_max_ was 0.67 ± 0.29 h, and MRT_0-∞_ was 6.17 ± 2.01 h, suggesting a favorable PK profile. Additional parameters such as T_max_, clearance (CL), and elimination rate (ER) remained relatively consistent across the tested doses. The volume of distribution, V_z_, after oral drug delivery of 2 to 50 mg/kg of **LG157**, reached 3.57 ± 1.35 to 6.43 ± 1.12 L/Kg, respectively, consistent with the wide tissue distribution detected in **Figure 3**. The standard deviation in each PK parameter among the three rats used in this study was less than 50%, indicating that the individual variance of these PK properties was small in rats. Comparative data from mice and rats suggested that effective therapeutic exposure levels could also be achieved in rats.^5^ Overall, these findings strongly indicate that **LG157** has a promising PK profile, making it a suitable candidate for further therapeutic evaluation.

**Figure 3.**
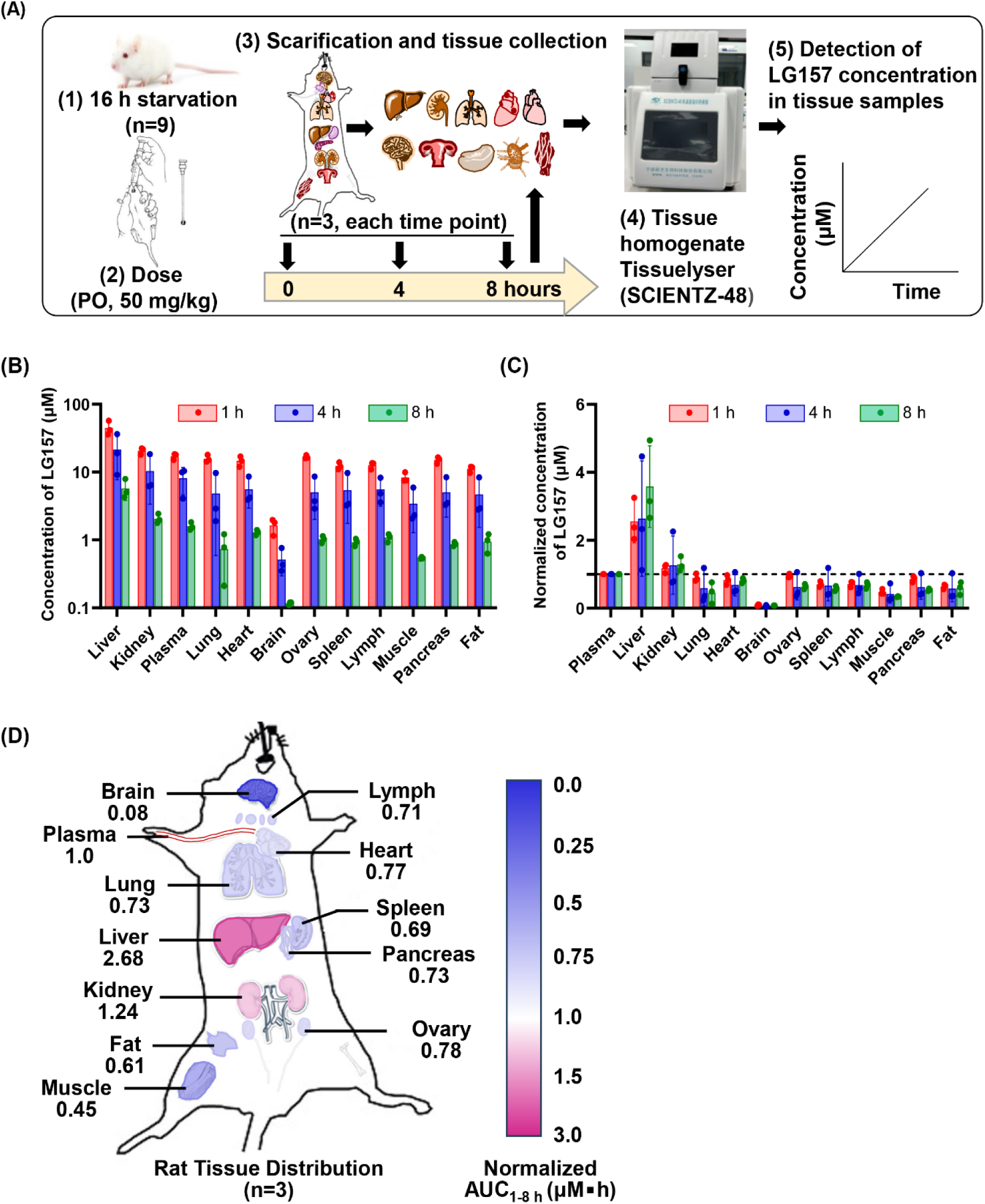
Tissue distribution of **LG157**. Wistar rats received a single oral dose of 50 mg/kg **LG157** and three female Wistar rats, 200-230 g each, were used at each time point. The peripheral blood and different tissues were collected at 1, 4, or 8 h after drug administration. The plasma drug concentration was determined by LC-MS/MS. (A) Flow chart of the tissue collection scheme for drug tissue distribution studies. (B) **LG157** concentrations in different tissues at the indicated time points after administration. (C) **LG157** concentrations in tissues after normalized by the plasma drug concentration. (D) AUC_1-8 h_ (μM▪h) of **LG157** in different tissues normalized by the plasma drug concentration.

**Table 3.**
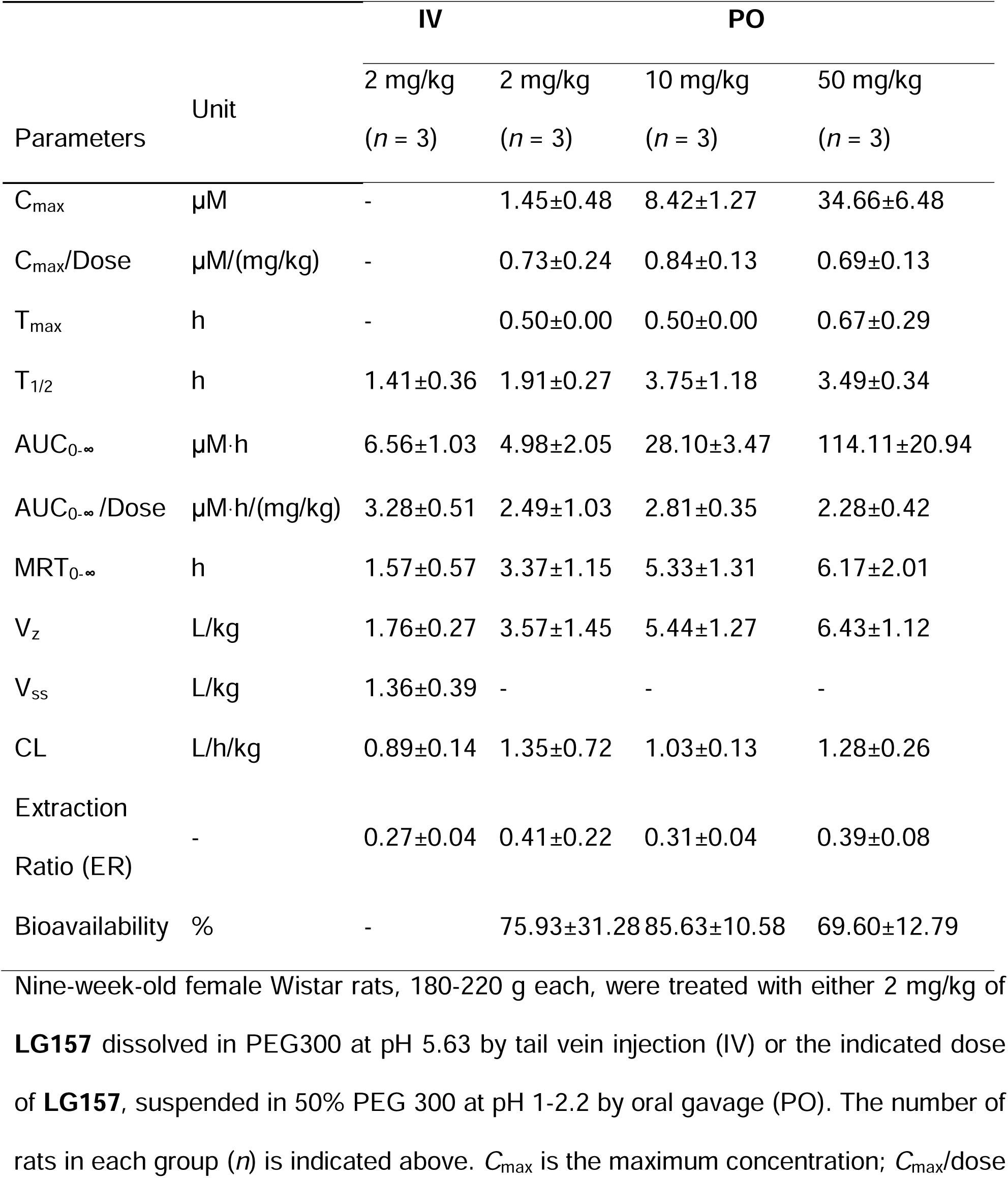
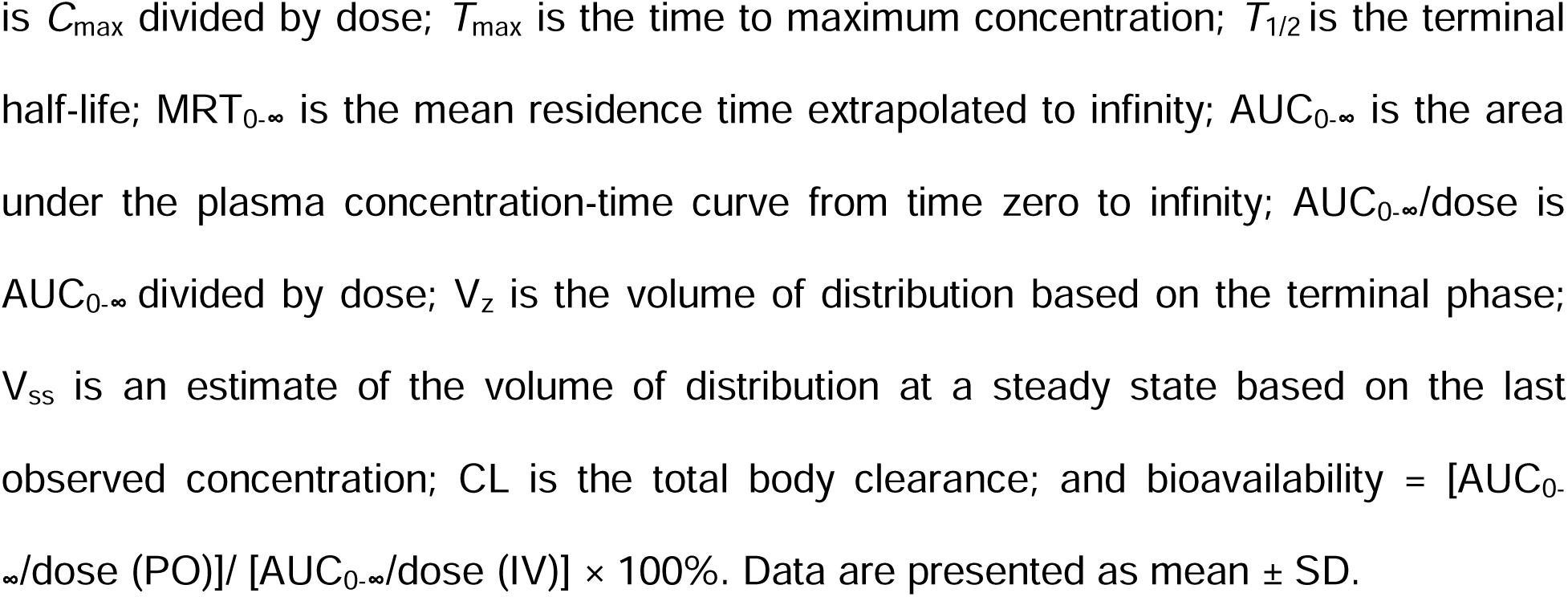
Pharmacokinetic parameters of LG157 after a single oral or intravenous administration to Wistar rats.

| Parameters | Unit | IV | PO |  |  |
| --- | --- | --- | --- | --- | --- |
|  |  | 2 mg/kg<br>(n = 3) | 2 mg/kg<br>(n = 3) | 10 mg/kg<br>(n = 3) | 50 mg/kg<br>(n = 3) |
| C<sub>max</sub> | μM | - | 1.45±0.48 | 8.42±1.27 | 34.66±6.48 |
| C<sub>max</sub>/Dose | μM/(mg/kg) | - | 0.73±0.24 | 0.84±0.13 | 0.69±0.13 |
| T<sub>max</sub> | h | - | 0.50±0.00 | 0.50±0.00 | 0.67±0.29 |
| T<sub>1/2</sub> | h | 1.41±0.36 | 1.91±0.27 | 3.75±1.18 | 3.49±0.34 |
| AUC<sub>0-∞</sub> | μM·h | 6.56±1.03 | 4.98±2.05 | 28.10±3.47 | 114.11±20.94 |
| AUC<sub>0-∞</sub> /Dose | μM·h/(mg/kg) | 3.28±0.51 | 2.49±1.03 | 2.81±0.35 | 2.28±0.42 |
| MRT<sub>0-∞</sub> | h | 1.57±0.57 | 3.37±1.15 | 5.33±1.31 | 6.17±2.01 |
| V<sub>z</sub> | L/kg | 1.76±0.27 | 3.57±1.45 | 5.44±1.27 | 6.43±1.12 |
| V<sub>ss</sub> | L/kg | 1.36±0.39 | - | - | - |
| CL | L/h/kg | 0.89±0.14 | 1.35±0.72 | 1.03±0.13 | 1.28±0.26 |
| Extraction Ratio (ER) | - | 0.27±0.04 | 0.41±0.22 | 0.31±0.04 | 0.39±0.08 |
| Bioavailability | % | - | 75.93±31.28 | 85.63±10.58 | 69.60±12.79 |

Nine-week-old female Wistar rats, 180-220 g each, were treated with either 2 mg/kg of **LG157** dissolved in PEG300 at pH 5.63 by tail vein injection (IV) or the indicated dose of **LG157**, suspended in 50% PEG 300 at pH 1-2.2 by oral gavage (PO). The number of rats in each group (n) is indicated above. C<sub>max</sub> is the maximum concentration; C<sub>max</sub>/dose
is $C_{\max}$ divided by dose; $T_{\max}$ is the time to maximum concentration; $T_{1/2}$ is the terminal half-life; $MRT_{0-\infty}$ is the mean residence time extrapolated to infinity; $AUC_{0-\infty}$ is the area under the plasma concentration-time curve from time zero to infinity; $AUC_{0-\infty}/\text{dose}$ is $AUC_{0-\infty}$ divided by dose; $V_z$ is the volume of distribution based on the terminal phase; $V_{ss}$ is an estimate of the volume of distribution at a steady state based on the last observed concentration; CL is the total body clearance; and bioavailability = $[AUC_{0-\infty}/\text{dose (PO)}] / [AUC_{0-\infty}/\text{dose (IV)}] \times 100\%$ . Data are presented as mean ± SD.

### Wide tissue distribution of LG157

Our investigation into the tissue and organ distribution of **LG157** leveraged the PEG 300 formulation. As illustrated in **Figure 3**, the widespread distribution of **LG157** indicates the ability of this compound to traverse various biological barriers, a favorable attribute for oncology drugs. Our findings indicated the highest deposition of **LG157** in the liver, as quantified by an average AUC_1-8h_ value of 153.70 μM▪h. Conversely, the brain showed the lowest uptake of the drug, with an average AUC_1-8h_ value of 4.49 μM▪h (**Figure 3D**). This discrepancy in tissue deposition suggests that **LG157** could accumulate in the liver where its drug-binding proteins may reside, or it is a substrate for a transporter protein that’s highly expressed in the liver. Certain drugs naturally gravitate toward the liver due to its high vascularization and its role in detoxification and metabolism.^35,36^ **LG157** may simply follow this common biodistribution pattern. An alternative explanation for the unexpectedly elevated liver concentrations could be the potential inability of liver-specific CYP enzymes to metabolize **LG157** effectively. Given that the liver is a critical site for drug metabolism, any resistance to such processes could lead to the accumulation of **LG157** within the liver.^35^

### Development of SEDDS for LG157

**LG157** falls into class I of BCS, possessing favorable characteristics such as good solubility, good permeability, and decent oral bioavailability in the preclinical formulation. These features make it a promising candidate for conventional formulation approaches. However, we decided to explore its potential to be formulated as SEDDS to take advantage of its unique features of SEDDS. For this purpose, we first tested the solubility of this compound in a panel of excipients including oils, surfactants cosolvents/cosurfactants, and polymers. The oil, surfactant, and cosurfactant with the maximal solubility for **LG157** were identified, with the solubility of **LG157** in Oleic acid at 135.61 mg/ml, surfactant PEG400 at 147.91 mg/ml, and cosurfactants Tween-20 at 79.55 mg/ml (Figures 4).

**Figure 4.**
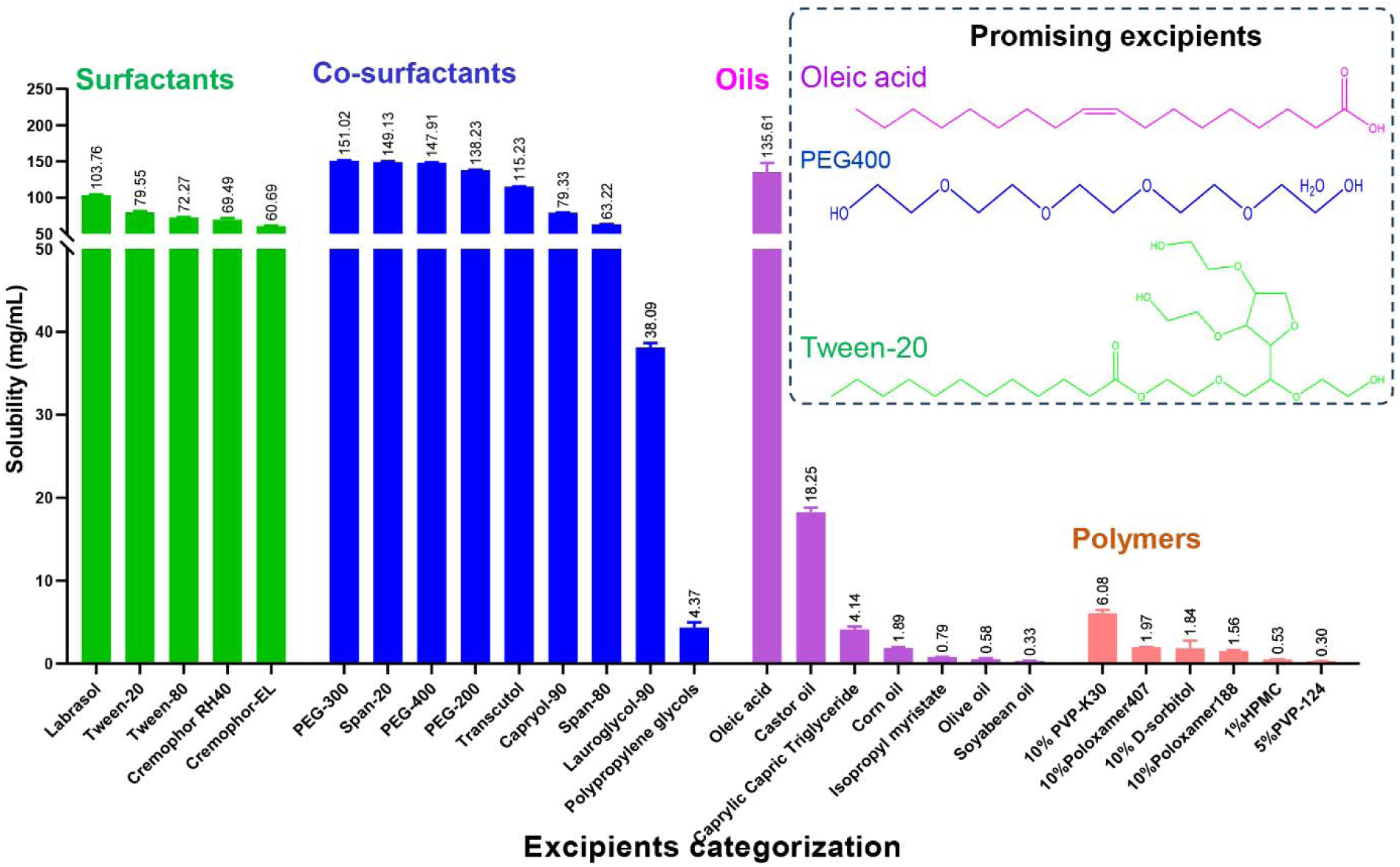
Solubility of **LG157** in various excipients. The solubility of **LG157** was determined after incubation of **LG157** powder with the indicated excipient at 37°C for 72 h. The boxed chemicals were used for SEDDS studies.

A ternary phase diagram represents all possible mixture ratios of three optimal solvents. The dotted areas illustrate micro emulsification regions along with the highest and lowest probability of forming microemulsions ^22^ (Figure 5). A SEDDS was designed and optimized through the design of an expert (DOE) to determine the optimal ratios of excipients. Thirteen sets of experiments were generated with the help of DOE, demonstrating a drug loading capacity of 140.6 to 200.5 mg of **LG157** in 1 ml of SEDDS (**Table 4).**

**Figure 5.**
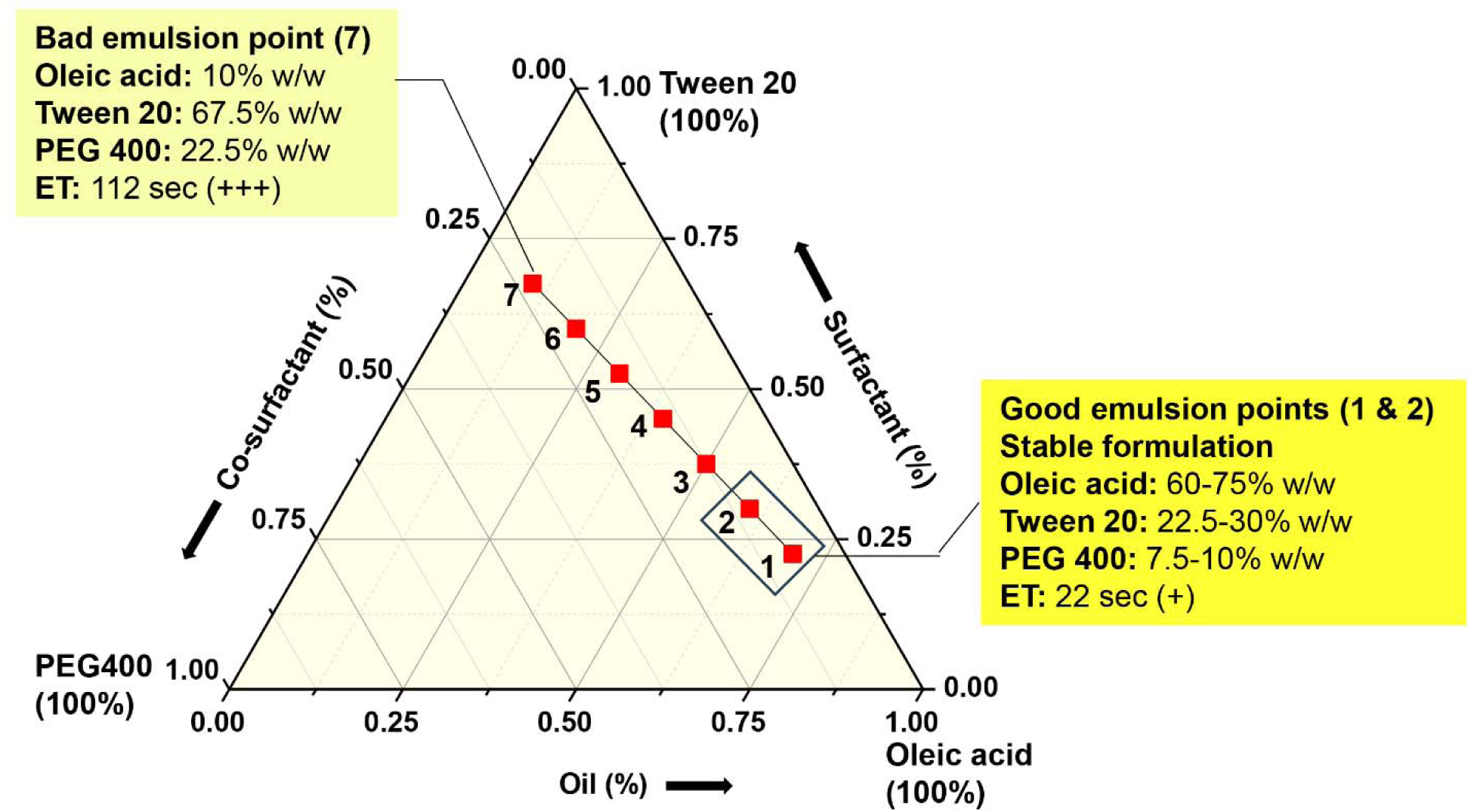
Ternary phase diagram for a mixture of Oil (Oleic acid), Surfactant (Tween-20), and Co-surfactant (PEG400). S_mix_ ratio (3:1) and Oil: S_mix_ diverse weight ratios (% w/w), including 7:3, 6:4, 5:5, 4:6, 3:7, 2:8, and 1:9. The red points numbered 1 to 7 indicate the varying percentage of excipients used in formulation within the emulsification range. The plus sign indicates the emulsification time (ET) range for the indicated 7 points as follows: Points 1 and 2: + (lowest), Points 3-6: ++ (medium), and Point 7: +++ (highest).

**Table 4.** Independent variables (X1, X2) and Experimental Design Matrix of Central Composite Design (CCD) Design for LG-157 for Drug Loading Response (Y1)

| Experiments | Variables |  | Responses |
| --- | --- | --- | --- |
|  | X1<br>(% of oil) | X2<br>(% of Smix) | Drug<br>loading (Y1)<br>mg/ml |
| 1 | 65 | 3 | 193.9 |
| 2 | 42.5 | 2.5 | 180.1 |
| 3 | 42.5 | 2.5 | 179.9 |
| 4 | 65 | 2 | 198.2 |
| 5 | 42.5 | 2.5 | 177.4 |
| 6 | 20 | 2 | 148.3 |
| 7 | 10.6802 | 2.5 | 140.6 |
| 8 | 42.5 | 1.79289 | 173.1 |
| 9 | 42.5 | 2.5 | 172.0 |
| 10 | 42.5 | 3.20711 | 163.3 |
| 11 | 42.5 | 2.5 | 182.8 |
| 12 | 74.3198 | 2.5 | 200.5 |
| 13 | 20 | 3 | 158.4 |

The three-dimensional (3D) response surfaces and contour plots (2D) for drug loading (Y_1_) are displayed in Figure 6. Two factors, X_1_: (A: percentage of oil, %); and X2: (B: Smix: ratio of surfactant: co-surfactant) along with their interaction had a valid influence on response Y_1._ Upon the addition of more oil and reduction of the Smix (surfactant: co-surfactant ratio) amount, the drug loading increased gradually. The response surface plot and contour plot show that the combination of surfactant, co-surfactant, and oil ratios influence drug loading. The ANOVA for drug loading is depicted in **Table S3**. From the table, the model is significant *(*P*<0.001), but the lack-of-fit terms were not, anyway. The model F-value of 31.15 implies the model is statistically significant. There is only a 0.01% chance that an F-value this large could occur due to noise. The Lack of Fit F-value of 2.11 implies the Lack of Fit is not significant relative to the pure error. The *p*-value evaluates the significance of each model for drug loading. The coefficient of determination (*R*^2^) of the model **LG157**-drug loading (Y_1_) was 0.957 and CV% less than 10.

**Figure 6.**
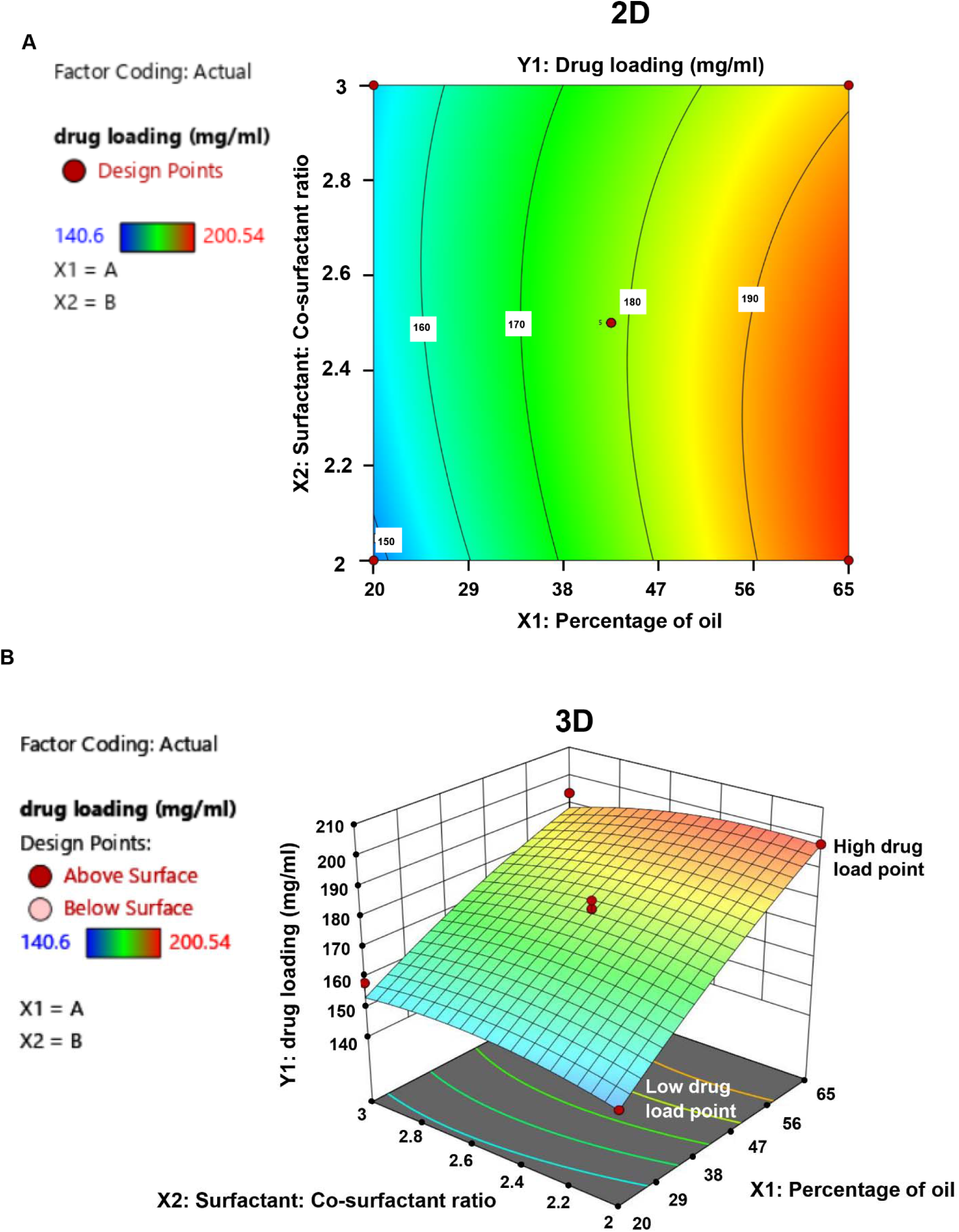
Design expert plot. (A) Two-dimensional (2D) contour plot and (B) the three-dimensional (3D) response surface plot show the effects of independent factors on drug loading. Red points on 2D and 3D plots represent different combination levels of oil percentage and S_mix_ that result in the desired response for drug loading.

**Figure 7.**
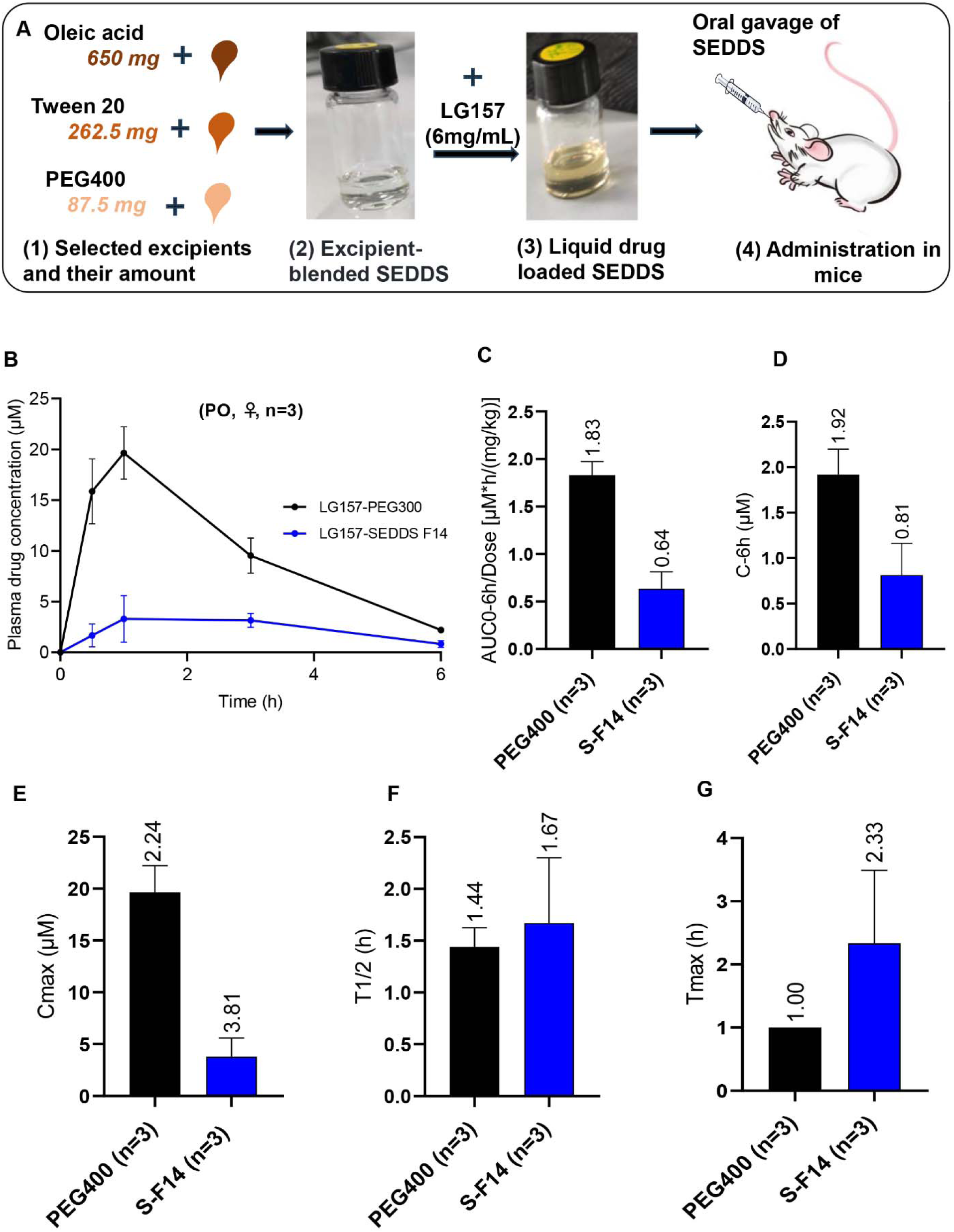
Pharmacokinetic parameters of **LG157** in SEDDS-F14. **LG157** was formulated in PEG300 or SEDDS-F14 and administered at a dose of 30 mg/kg to female BALB/c mice by oral gavage (PO). n: the number of mice used in each group. (A) Schematic representation of the liquid-SEDDS-F14 preparation procedure. (B) Plasma drug concentration-time profile of **LG157** in SEDDS-F14 and PEG300. (C) The area under the plasma drug concentration-time curve from time zero to 6 h (AUC_0-6h_) divided by dose. (D) Plasma drug concentration at 6h (C_6h_). (E) The maximum drug concentration in plasma (*C*_max_). (F) Half-life time (t_1/2_). (G) Time of peak plasma drug concentration (T_max_).

The results of the 13 sets of formulation experiments suggest excipient composition for SEDDS-F14 and predicted its drug loading capacity. The final formulation, SEDDS-F14 was generated from the DOE with a desirability of 1, indicating that the desired target response for a drug loading of 188.7 mg/ml has been achieved (**Table S4**) It meets the requirements for the projected one-pill-a-day clinical use of 150 mg of **LG157**.

### PK properties of LG157-SEDDS-F14

For PK analysis, SEDDS-F14 was loaded with 6 mg/ml of **LG157**, equal to 30 mg/kg body weight for mice, and proceeded for analysis. The formulation composition and other characterized details of SEDDS-F14 are shown in **Table S5.** PK analysis represented the acquired plasma concentration *vs*. time profiles after the oral administration of a single dose of SEDDS-F14 to male rats (Figure 5). AUC_0-6 h_ was 0.64 [µM*h/(mg/kg)], this plasma drug exposure was 2.55-fold lower than that of the PEG 400 formulation but was still much higher than the threshold of 0.1. SEDDS-F14 showed a slight improvement in half-life value, and the time to reach the maximum drug concentration in plasma (*T*_max_) was 2.33 h, longer than that of the PEG 400 formulation. Adequate drug exposure to **LG157** by using SEDDS opens a new avenue to test the therapeutic efficacy of this compound and further modify this formulation for improved efficacy and reduced toxicity.

## DISCUSSIONS AND CONCLUSION

Our study has revealed the promising PK profile of **LG157**, highlighting its desirable druglike properties, wide tissue distribution, high stability under various stress conditions, and favorable solubility characteristics both in aqueous media and various excipients used in SEDDS. The synthesis of **LG157** is straightforward and feasible in 3 steps. ^5^ These experimental findings, coupled with computational modeling data, strongly support the developability of **LG157** as an orally administered therapeutic agent.

We have successfully developed a preliminary **LG157**-SEDDS formulation, setting the stage for further optimization efforts. Our primary goal moving forward is to refine the **LG157**-SEDDS to enhance **LG157**’s bioavailability and modify its PK profile. Specifically, we aim to achieve a controlled release mechanism to mitigate peak plasma drug concentration (C_max_) which is often associated with cardiac toxicity, while simultaneously extending the duration of effective plasma drug levels for sustained target inhibition.

Optimization strategies for SEDDS will focus on selecting appropriate excipients, and fine-tuning release profiles. Currently, our SEDDS formulation has been developed analytically, utilizing a combination of trial-and-error methods, component ratio adjustments, ternary phase diagrams, and response surface methodology (RSM), However, these methods, while effective, are often labor-intensive and require exploration of numerous combinations. To streamline the development process, future efforts will implement an integrated approach combining a trial of other Smix ratios, alter excipients combination, and Hydrophilic-Lipophilic Balance (HLB) with RSM^37^, along with transitioning from liquid to solid SEDDS. This approach is expected to reduce the number of experimental trials needed to optimize the SEDDS for achieving the desired **LG157** drug release profile.

The encouraging results and advantages of the SEDDS approach, as demonstrated in previous studies^10,11^, compared to conventional formulations, provide a strong impetus to continue this research. The conventional understanding of SEDDS primarily revolves around its bioavailability enhancement and inhibition of multidrug resistance pumps. Conceptually, SEDDS could also include a nontoxic hydrophobic anti-cancer agent with a mechanism of action complementing that of **LG157**. *Trans* vaccenic acid (TVA) is a naturally occurring fatty acid that modulates natural killer cell function and elicits anti-tumor immunity.^38^ Incorporating TVA or another hydrophobic compound with a similar activity into **LG157**-SEDDS formulation may enhance the efficacy of **LG157** in treating tumors. The potential integration of immunity-triggering hydrophobic compounds into SEDDS opens new horizons, providing a foundation for holistic strategies beyond traditional bioavailability considerations.

## ASSOCIATED CONTENT

### Supporting information

The supporting information file comprises supporting Tables (**Table S1-S5**). **Notes**

The authors declare no competing financial interest.

### Funding source

The J. Michael Bishop Institute of Cancer Research receives funding through an endowment from Anticancer Bioscience, a company actively engaged in the commercial development of cancer therapeutics.

## ABBREVIATIONS

CCD-RSM: Central Composite Design-Response Surface Methodology
DOE: Design of Expert
DMA: Dimethylacetamide
GIT: gastrointestinal tract
HPMC: Hydroxypropyl methylcellulose
PEG300: Polyethylene glycol 300
SEDDS: self-emulsifying drug delivery system
PK: pharmacokinetics

## ACKNOWLEDGMENTS

We are grateful to all divisions of Anticancer Bioscience for their critical review and constructive suggestions.

