## Supplementary information file for "Pharmacokinetics, Tissue Distribution, and Formulation Study of a Small-molecule Inhibitor of MKLP2, LG157"

*****Correspondence authors

**Table S1.** SwissADME Predictions of the Pharmacokinetics Properties for **LG157**

| **Boiled egg (Brain or Intestinal)** | | | **CYP-inhibitors** | | | | |
| --- | --- | --- | --- | --- | --- | --- | --- |
| GI absorption | BBB permeant | Pgp substrate | CYP1A2 | CYP2C19 | CYP2C9 | CYP2D6 | CYP3A4 |
| High | Yes | No | +++ | ++ | + | ++ | ++ |

| **Drug likeness** | | | | **Medicinal chemistry** | | | |
| --- | --- | --- | --- | --- | --- | --- | --- |
| Lipinski #violations | Ghose #violations | Veber #violations | Bioavailability Score | PAINS #alerts | Brenk #alerts | Lead likeness | Synthetic score |
| Accepted | Accepted | Accepted | 0.55 | 0 | 0 | Yes | 3.04 |

**Table S2**. In Silico Predictions of the Drug-Likeness and Medicinal Chemistry Parameters for **LG157**

**Table S3.** Analysis of Variance (ANOVA) of the Drug Loading of **LG157**

| **Source** | **Sum of Squares** | **df** | **Mean Square** | **F-value** | **p-value** |  |
| --- | --- | --- | --- | --- | --- | --- |
| **Model** | 3811.79 | 5 | 762.36 | 31.15 | 0.0001 | significant |
| A-A | 3619.22 | 1 | 3619.22 | 147.87 | < 0.0001 |  |
| B-B | 8.01 | 1 | 8.01 | 0.3274 | 0.5851 |  |
| AB | 52.49 | 1 | 52.49 | 2.14 | 0.1865 |  |
| A² | 47.66 | 1 | 47.66 | 1.95 | 0.2055 |  |
| B² | 100.19 | 1 | 100.19 | 4.09 | 0.0827 |  |
| **Residual** | 171.32 | 7 | 24.47 |  |  |  |
| Lack of Fit | 105.02 | 3 | 35.01 | 2.11 | 0.2415 | not significant |
| Pure Error | 66.31 | 4 | 16.58 |  |  |  |
| **Cor Total** | 3983.11 | 12 |  |  |  |  |

**Table S4**. Desirability Data for F14-SEDDS Desirability and Solution after 13 sets of DOE Experiments

| **Number** | **A** | **B** | **drug loading** | **Desirability** |  |
| --- | --- | --- | --- | --- | --- |
| **1** | **65.000** | **3.000** | **188.704** | **1.000** | **Selected** |
| 2 | 58.853 | 3.000 | 185.117 | 0.929 |  |
| 3 | 65.000 | 2.852 | 191.993 | 0.923 |  |
| 4 | 55.826 | 3.000 | 183.208 | 0.892 |  |

**Table S5: LG157**-SEDDS-F14 Formulations Composition, Percentage Transmittance, Emulsification time, and Lipid Formulation Classification System (LFCS)

| **Formulation composition SEDDS-F14** | | | **Characteristic details** | | | | | |
| --- | --- | --- | --- | --- | --- | --- | --- | --- |
| Oil (mg)  Oleic acid | Surfactant (mg)  Tween -20 | Co-surfactant (mg)  PEG-400 | Drug  load (mg) | Drug  load (%) | Transmittance%  (OD 620nm) | Emulsification time(sec) | Clarity | LFCS type |
| 650 | 262.5 | 87.5 | 6 | 0.6 | 97 | 26 | Transparent | II |
